# Calpain-2 mediates SARS-CoV-2 entry and represents a therapeutic target

**DOI:** 10.1101/2022.11.29.518418

**Authors:** Qiru Zeng, Avan Antia, Maritza Puray-Chavez, Sebla B. Kutluay, Siyuan Ding

## Abstract

Since the beginning of the coronavirus disease 2019 (COVID-19) pandemic, much effort has been dedicated to identifying effective antivirals against severe acute respiratory syndrome coronavirus 2 (SARS-CoV-2). A number of calpain inhibitors show excellent antiviral activities against SARS-CoV-2 by targeting the viral main protease (M^pro^), which plays an essential role in processing viral polyproteins. In this study, we found that calpain inhibitors potently inhibited the infection of a chimeric vesicular stomatitis virus (VSV) encoding the SARS-CoV-2 spike protein, but not M^pro^. In contrast, calpain inhibitors did not exhibit antiviral activities towards the wild-type VSV with its native glycoprotein. Genetic knockout of calpain-2 by CRISPR/Cas9 conferred resistance of the host cells to the chimeric VSV-SARS-CoV-2 virus and a clinical isolate of wild-type SARS-CoV-2. Mechanistically, calpain-2 facilitates SARS-CoV-2 spike protein-mediated cell attachment by positively regulating the cell surface levels of ACE2. These results highlight an M^pro^-independent pathway targeted by calpain inhibitors for efficient viral inhibition. We also identify calpain-2 as a novel host factor and a potential therapeutic target responsible for SARS-CoV-2 infection at the entry step.

## INTRODUCTION

High mutation rates of SARS-CoV-2 pose great challenges for antiviral drug development and treatment of COVID-19 patients. Thus far, most antiviral strategies have directly targeted key viral factors involved in the SARS-CoV-2 replication cycle [1]. Remdesivir (Gilead) and molnupiravir (Merck) represent two FDA authorized antiviral drugs that inhibit the SARS-CoV-2 RNA-dependent RNA polymerase (RdRp) [2]. In addition to RdRp, the viral main protease (M^pro^) has been a drug target of great interest due to its fundamental role in processing the viral polyproteins. In a series of studies, M^pro^ inhibitors, including Paxlovid (Pfizer), boceprevir, GC376, and various calpain inhibitors, were reported to potently supress SARS-CoV-2 replication in different cell types and in pre-clinical animal models [3, 4].

In a recent study from our group [5], we performed a drug repurposing screen and identified several compounds that potently block SARS-CoV-2 infection. One such compound is MG132, a commonly used 26S proteasome inhibitor. MG132 was previously reported to impair SARS-CoV replication by inhibiting the host cysteine protease m-calpain, also known as calpain-2 (encoded by *CAPN2*), thus functioning through a proteosome-independent pathway [6]. Several pieces of evidence led us to hypothesize that host calpain proteases may be required for SARS-CoV-2 infection. First, MG132 inhibited SARS-CoV-2 replication, while ubiquitin-activating enzyme E1 inhibitor PRY-41 and two other proteasome inhibitors bortezomib and lactacystin did not. Second, E-64, which inhibits endosomal cathepsins, papain, and calpain, inhibited SARS-CoV-2 more robustly than chloroquine, which only targets cathepsins and not calpain. Third, calpain inhibitor II and calpeptin suppressed SARS-CoV-2 replication [4, 7], although the mechanism was postulated to be mediated by interfering with activities of M^pro^.

Of note, in our follow-up studies described here, we found that MG132 also exhibited antiviral activities against a chimeric vesicular stomatitis virus (VSV) expressing SARS-CoV-2 spike protein (VSV-SARS-CoV-2) [8], but not expressing M^pro^. The lack of inhibition against wild-type (WT) VSV led us to hypothesize that (1) SARS-CoV-2 spike protein may be an additional viral target of calpain and protease inhibitors; and (2) calpain proteins themselves may be crucial host factors for SARS-CoV-2 infection. In this paper, we confirm these hypotheses and identify CAPN2 as a novel pro-viral host factor that aids in the entry of SARS-CoV-2. We demonstrate that the absence of *CAPN2* reduces viral binding to host cells and RNA production during early steps of the SARS-CoV-2 replication cycle. The findings provide mechanistic insights into the cellular process of SARS-CoV-2 entry and offer an additional explanation to the mechanism of action of calpain inhibitors.

## RESULTS

### MG132 preferentially inhibits the infection of VSV-SARS-CoV-2 but not VSV

In a recent antiviral compound screen that we conducted using a recombinant SARS-CoV-2 mNeonGreen reporter virus [5], multiple compounds efficaciously inhibited viral infection in Vero E6 cells. We validated 18 of the top hits using recombinant VSV eGFP reporter viruses that either encode the SARS-CoV-2 spike protein or the native VSV-G [8]. Among the 18 compounds that we tested, most showed a dose-dependent inhibition of VSV-SARS-CoV-2 and VSV infections in MA104 cells (**Fig. 1**). Nigericin, brefeldin A, and 3-isobutyl-1-methylxanthine (IBMX) had effective concentration to reach 50% inhibition (EC_50_) values lower than 2 μM against both VSV-SARS-CoV-2 and VSV infections (**Fig. 1**). Nitazoxanide was recently reported to inhibit SARS-CoV-2 infection [9] and we observed similar results (**Fig. S1**). In addition, we noticed that MG132, a broad-spectrum proteasome inhibitor, exhibited a 100-fold selectivity in antiviral activities against VSV (EC_50_ of 44.4 μM) and VSV-SARS-CoV-2 (EC_50_ of 0.64 μM) (**Fig. 1**). Notably, neither of the two reporter viruses express M^pro^ and the only difference lies in the VSV glycoprotein replaced by the SARS-CoV-2 spike protein. This data, together with the previous report of MG132 and SARS-CoV, led us to further test the antiviral activities of MG132 and other calpain inhibitors and their effect on the spike protein.

**Fig. 1.**
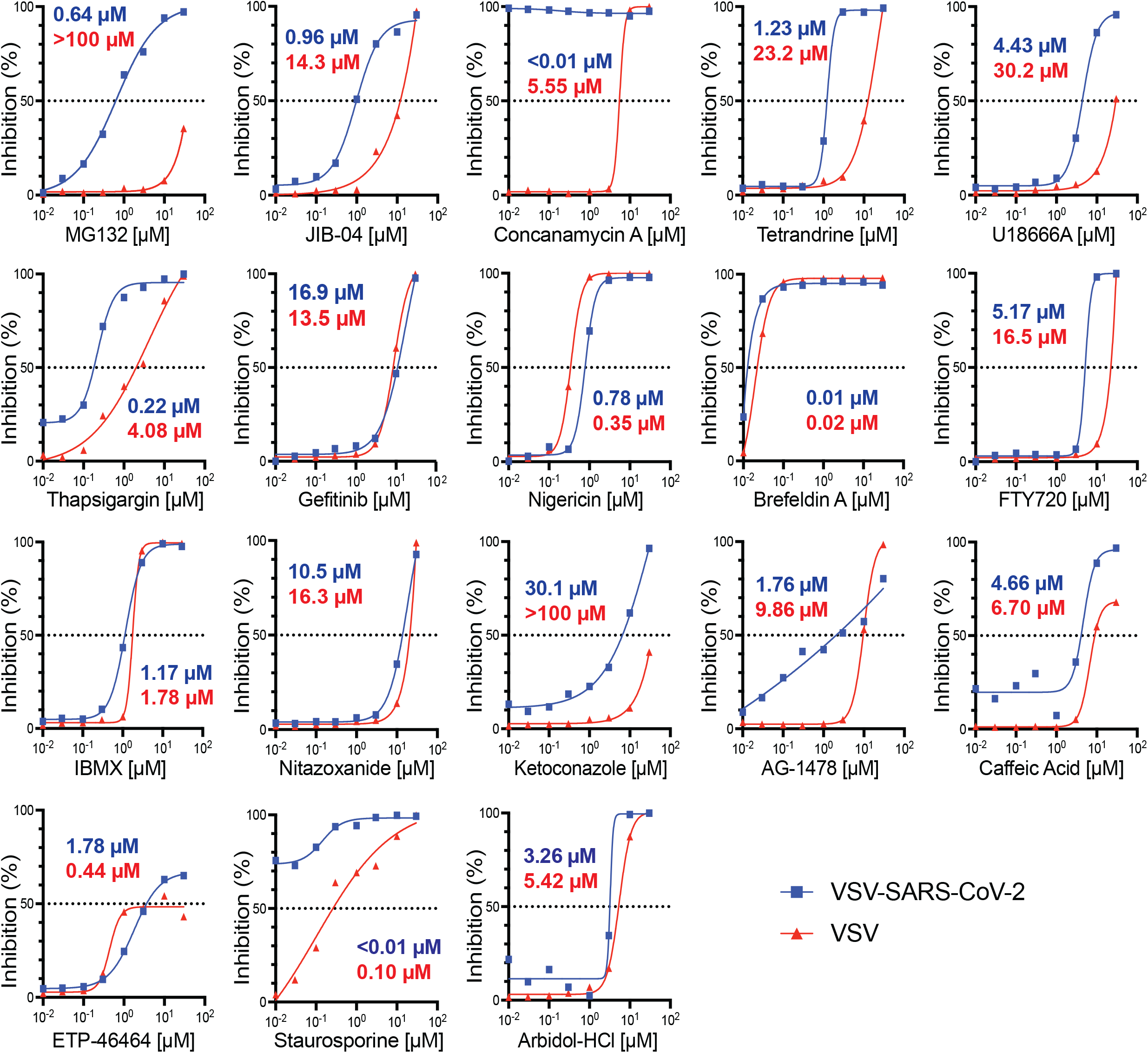
Small-molecule compounds inhibit VSV and VSV-SARS-CoV-2 infection. Screening of 18 compounds over a 24-hour infection period. MA104 cells were pre-treated with each compound for 1 hour at indicated concentrations ranging from 0.01 μM to 30 μM and then infected for 24 hours with either recombinant VSV-SARS-CoV-2 (MOI=1) or VSV (MOI=1). Quantified GFP signals are plotted as percentage of inhibition corresponding to dosage. EC_50_ values for each curve are indicated in blue (VSV-SARS-CoV-2) or red (VSV).

### Calpain inhibitors strongly inhibit VSV-SARS-CoV-2 infection

To examine whether MG132 targeting host calpain proteases accounts for the inhibition observed, we tested a set of commercially available calpain inhibitors, including ALLN (also known as MG101 or calpain inhibitor I), calpain inhibitor III, calpeptin, and E-64d, since these calpain inhibitors vary in their specificities targeting different members of the calpain family [10]. With the exception of calpeptin, none of the inhibitors were cytotoxic even at the highest concentration tested (**Fig. 2A**). All four calpain inhibitors exhibited potent inhibition against VSV-SARS-CoV-2 with EC_50_ values lower than 1.5 μM (**Fig. 2B**).

**Fig. 2.**
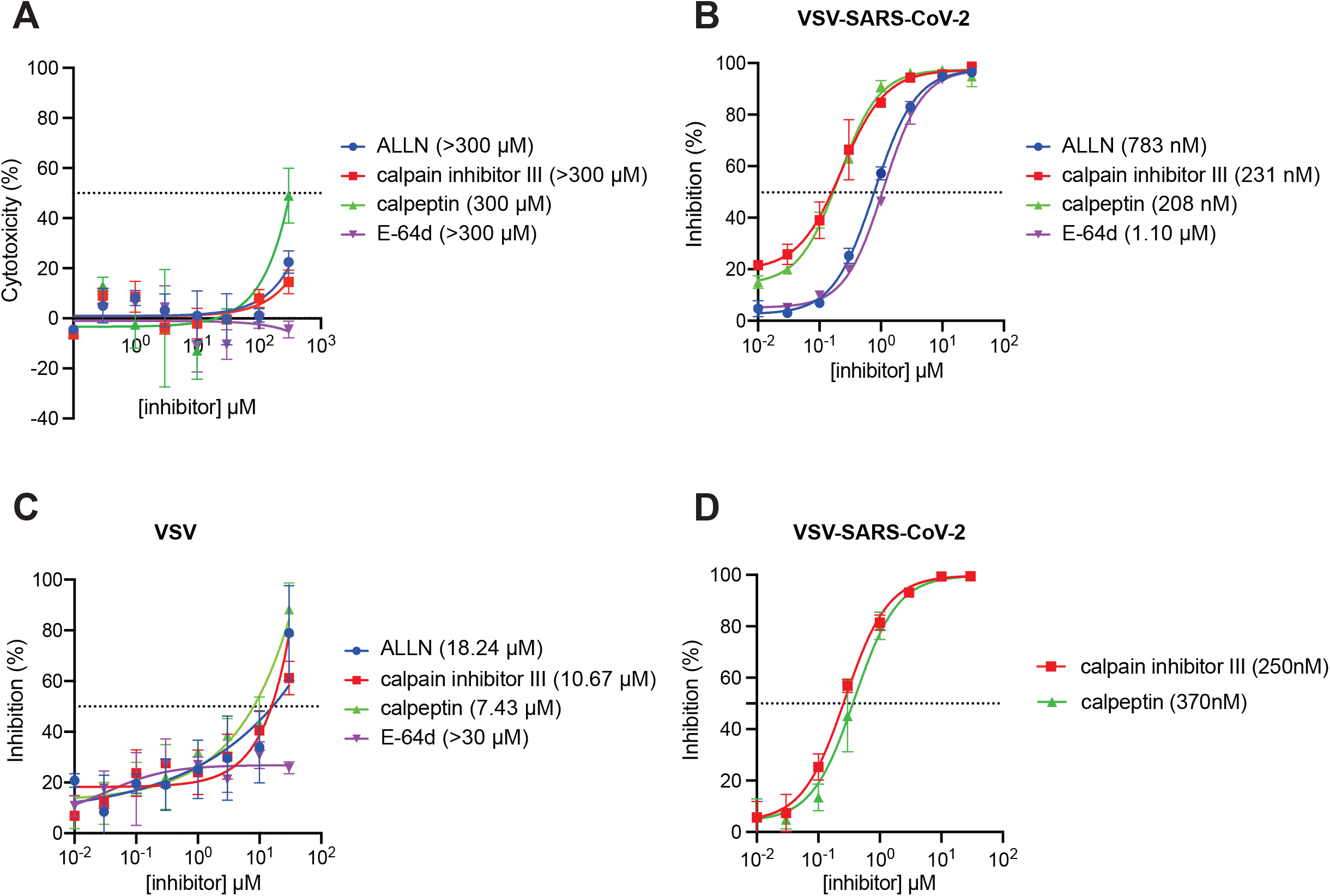
Calpain inhibitors potently inhibit VSV-SARS-CoV-2 infection. (A) Cytotoxicity assay of calpain inhibitors. MA104 cells were treated with ALLN, calpain inhibitor III, calpeptin, and E-64d for 25 hours and tested for cell viability. Percent cytotoxicity was plotted corresponding to dosage. CC_50_ values are as indicated. (B) MA104 cells were pretreated with ALLN, calpain inhibitor III, calpeptin, and E-64d at concentrations ranging from 0.01 μM to 30 μM for 1 hour prior to a 24-hour infection by VSV-SARS-CoV-2 (MOI=1). GFP signals were quantified and plotted as percentage of inhibition corresponding to dosage. EC50s values are as indicated. (C) Same as (B) except that VSV was used for infection instead of VSV-SARS-CoV-2. (D) Same as (B) except that Vero E6 cells were used instead of MA104 cells.

Calpain inhibitor III and calpeptin, which are more inhibitory against CAPN2 [10], showed stronger efficacy than ALLN and E-64d, with EC_50_ values of 231 nM and 208 nM, respectively (**Fig. 2B**). VSV was reported to not be sensitive to E-64d treatment [11]. Consistently, all four calpain inhibitors were substantially less inhibitory against VSV infection (**Fig. 2C**). For instance, calpain inhibitor III and calpeptin had EC_50_ values of 10.67 μM and 7.43 μM, respectively, indicating a more than 30-fold increase when compared to VSV-SARS-CoV-2 (**Fig. 2C**). In addition to MA104 cells, the antiviral activities of calpain inhibitor III and calpeptin against VSV-SARS-CoV-2 were confirmed in Vero E6 cells (**Fig. 2D**). Notably, VSV-SARS-CoV-2 does not encode M^pro^, the key therapeutic target identified in many previous protease inhibitor studies. Therefore, the contrasting results of inhibition of VSV-SARS-CoV-2 compared to VSV and the more pronounced inhibitory effects seen with calpain inhibitor III and calpeptin prompted us to hypothesize that these calpain inhibitors may play a role in interfering with the activities of the SARS-CoV-2 spike protein by inhibiting the host gene *CAPN2*.

### VSV-SARS-CoV-2 infection is significantly reduced in *CAPN2* knockout cells

To directly investigate the role of the host gene *CAPN2* in SARS-CoV-2 infection, we genetically knocked out *CAPN2* by lentivirus-mediated CRISPR/Cas9 in MA104 cells, which express the endogenous ACE2 receptor that is necessary for virus entry. The *CAPN2* knockout (KO) efficiency was validated by western blot (**Fig. 3A**). WT and *CAPN2* KO cells were infected with VSV-SARS-CoV-2 at different multiplicities of infection (MOIs) and infectivity was determined at different time points post infection. We found that VSV-SARS-CoV-2 replication, reflected by GFP signals, was highly attenuated in *CAPN2* KO cells under all conditions (**Fig. S2A**). Consistently, intracellular viral mRNA levels were reduced by approximately 4-fold in the absence of CAPN2 (**Fig. S2B**), suggesting a pro-viral role of CAPN2 in VSV-SARS-CoV-2 infection. To further corroborate our findings, we performed standard plaque assays of VSV-SARS-CoV-2 infections at MOIs of 1, 0.1, 0.01, and 0.001 in WT and *CAPN2* KO cells. 10 plaques from each group were selected and the diameters of plaque sizes were quantified. The plaques of VSV-SARS-CoV-2 formed in the KO cells had an average of diameter of 1 mm, significantly smaller than the 2 mm observed in the WT cells (**Fig. 3B**). To test whether this phenotype was associated with the spike protein, we performed similar plaque assays using VSV. No significant difference in sizes of VSV plaques was observed between WT and KO cells (**Fig. 3C**), suggesting that CAPN2 promotes the replication of VSV-SARS-CoV-2 but not VSV by acting on the spike protein or facilitating spike protein related functions.

**Fig. 3.**
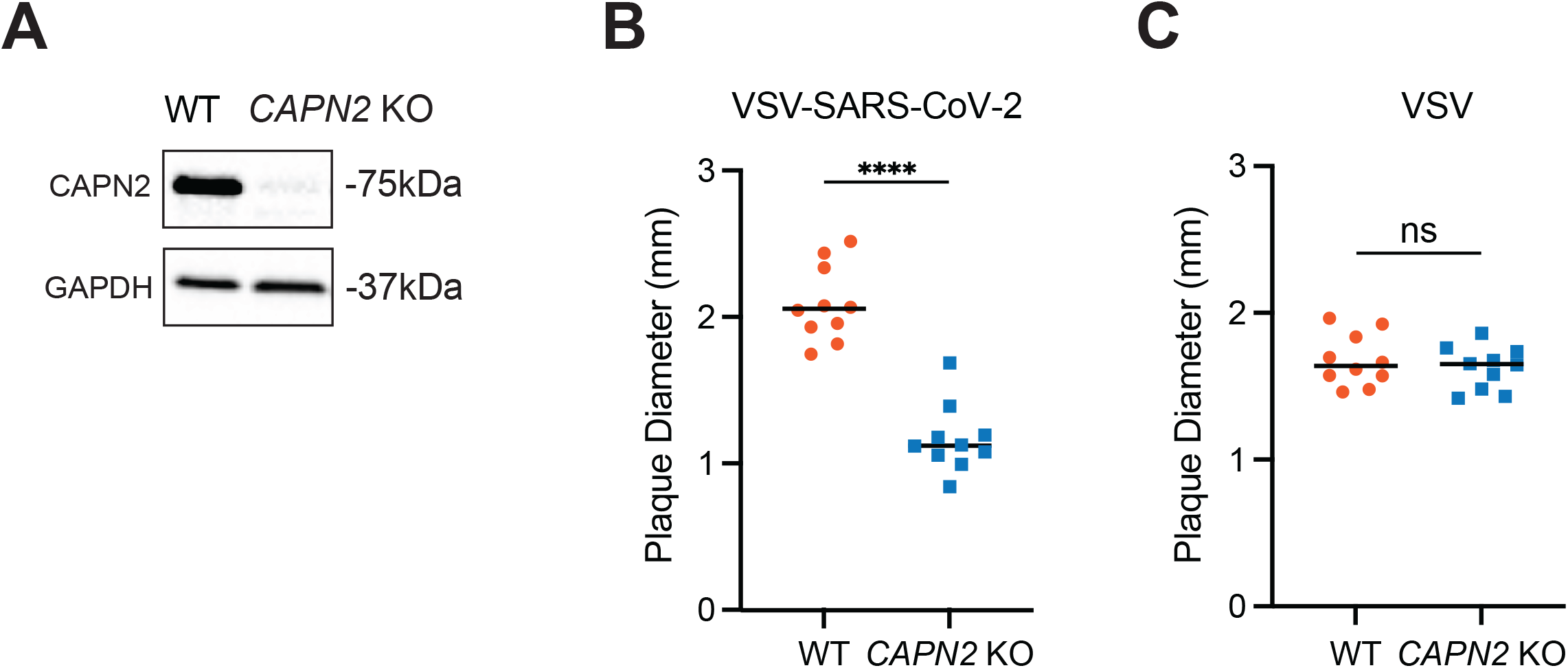
*CAPN2* KO cells have reduced VSV-SARS-CoV-2 infection. (A) Cell lysates of WT and *CAPN2* KO MA104 cells were harvested and the protein levels of CAPN2 and GAPDH were measured by western blot. (B) Plaque assays of VSV-SARS-CoV-2 were performed in MA104 cells. Images were taken at 72 hpi when clear-shaped plaques were observed. 10 plaques from each sample were selected and measured using microscopy. (C) Same as (B) except VSV was used instead and images were taken at 48 hpi.

### CAPN2 is required for an early step of the SARS-CoV-2 replication cycle

GFP signals from VSV-SARS-CoV-2 were reduced in *CAPN2* KO cells at 6 hours post infection (**Fig. S2A**), suggesting that CAPN2 functions to aid viral infection within a single replication cycle. We next sought to pinpoint the time point when CAPN2 exerts its pro-viral effect. A time-course experiment was performed by infecting WT and KO cells with VSV-SARS-CoV-2 and examining intracellular viral RNA levels at 1-6 hours post infection by RT-qPCR. Our results show that as early as 1 hour post infection, significantly lower viral mRNA levels were observed in the KO cells than those in the WT cells (**Fig. 4A**). Similar reduction was seen throughout the course of infection (**Fig. 4A**). A similar trend was reflected by the SARS-CoV-2 spike protein levels as the nascent protein synthesis was first visible starting at 4 hours post infection in WT cells, whereas it was barely detectable at 6 hours post infection in *CAPN2* KO cells (**Fig. 4B**).

**Fig. 4.**
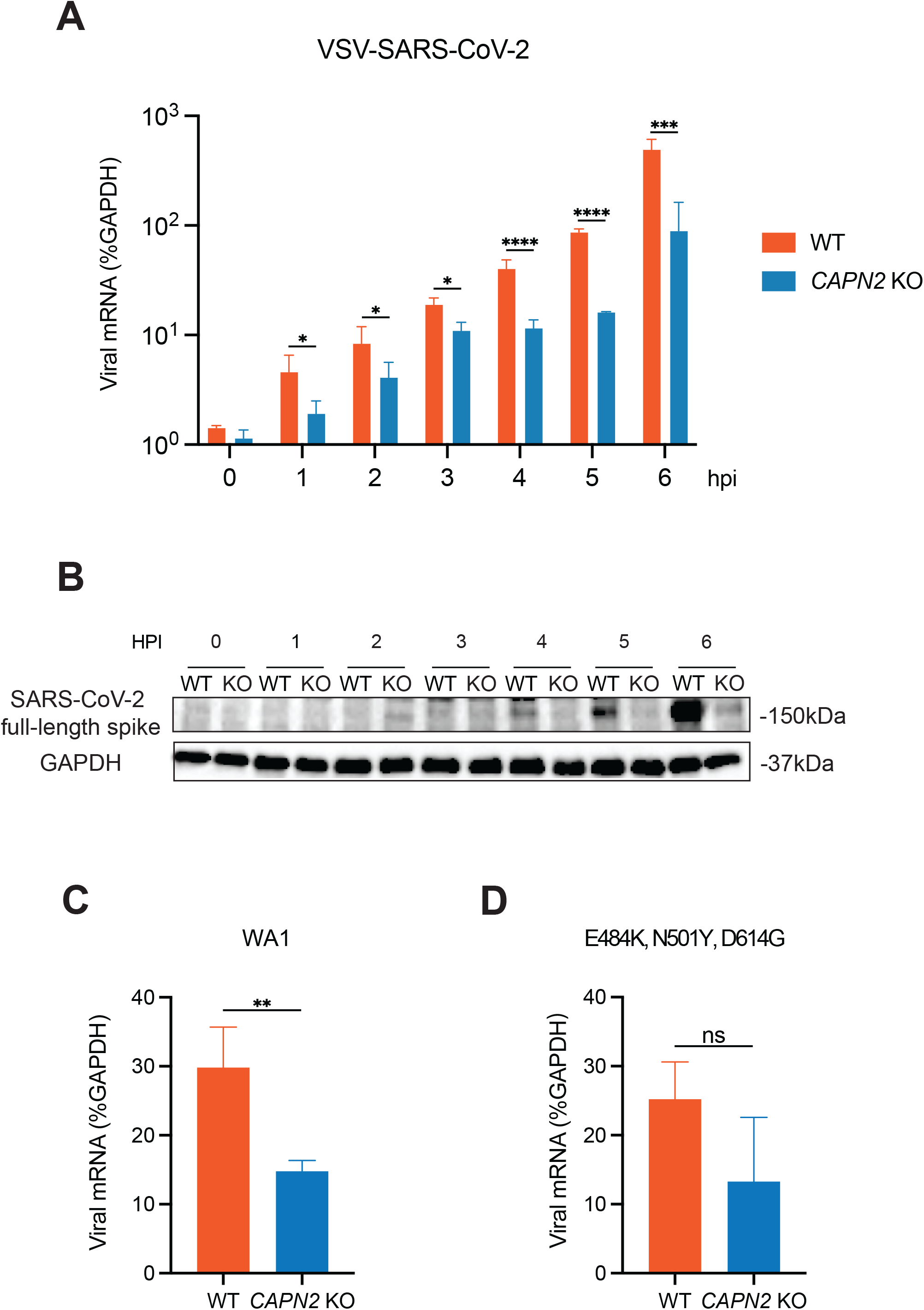
*CAPN2* deletion reduces SARS-CoV-2 infection. (A) Viral mRNA production at early time points post infection by VSV-SARS-CoV-2 (MOI=1) in WT and *CAPN2* KO MA104 cells. Cells were harvested at 0, 1, 2, 3, 4, 5, 6 hours post infection for RNA extraction followed by RT-qPCR analysis. Viral mRNA levels are shown relative to those of GAPDH. (B) Same as (A) except that SARS-CoV-2 full-length spike levels were measured by western blot instead. (C) Viral mRNA levels at 6 hours post infection by SARS-CoV-2 WA1 strain (MOI=0.1). Infected cells were measured for SARS-CoV-2 viral mRNA levels by qRT-PCR relative to those of GAPDH. (D) Same as (C) except a SARS-CoV-2 triple spike mutant strain was used instead.

Next, we set out to evaluate the early pro-viral role of CAPN2 in the context of a clinical isolate of infectious (?) SARS-CoV-2 (2019-nCoV/USA-WA1/2020 strain). Consistent with our findings with VSV-SARS-CoV-2, WT SARS-CoV-2 infection yielded lower intracellular viral mRNA levels in *CAPN2* KO cells than WT cells at 6 hours post infection (**Fig. 4C**). Concurrent SARS-CoV-2 variants accumulate multiple mutations in the spike protein that result in enhanced transmission and antibody evasion [12]. To that end, we tested a recombinant SARS-CoV-2 strain with spike mutations in three key residues E484K, N501Y, and D614G [13]. Interestingly, although the mRNA levels trended lower in *CAPN2* KO cells, the difference was not statistically significant (**Fig. 4D**), suggesting that the effect of CAPN2 on SARS-CoV-2 is potentially dependent on the nature of spike proteins.

### CAPN2 promotes SARS-CoV-2 binding to host cells

As suggested by the reduced levels of viral mRNA levels as early as 1 hpi (**Fig. 4A**), we reasoned that CAPN2 may play an important role at a very early time point of the SARS-CoV-2 replication cycle, i.e., virus binding and entry, which spike protein mediates. To test this hypothesis, we generated single clonal *CAPN2* KO MA104 cells, which were confirmed by Sanger sequencing (**Fig. S3A**), and took advantage of a classical cold binding assay [14] using VSV-SARS-CoV-2 to assess whether viral adsorption is negatively impacted by the lack of CAPN2. The assay was performed at 4 °C to allow virus binding to host cells but limit the energy required for virus endocytosis to gain entry into cells, followed by extensive wash and RT-qPCR analysis. As a positive control, we included a neutralizing antibody (2B04) that targets spike from the ancestral SARS-CoV-2 WA1 strain [15], the preincubation of which significantly reduced binding of the virus (**Fig. 5A**). Importantly, we found that the viral RNA levels from the virions bound to the KO cells were comparable to those in WT cells in the presence of antibody incubation and significantly lower than those in the WT cells (**Fig. 5A**). Similar binding defects in the KO cells were observed with WT SARS-CoV-2 WA1 strain (**Fig. 5B**). Viral binding in the KO cells were essentially reduced to the background levels similar to the antibody incubation controls (**Fig. 5B**).

**Fig. 5.**
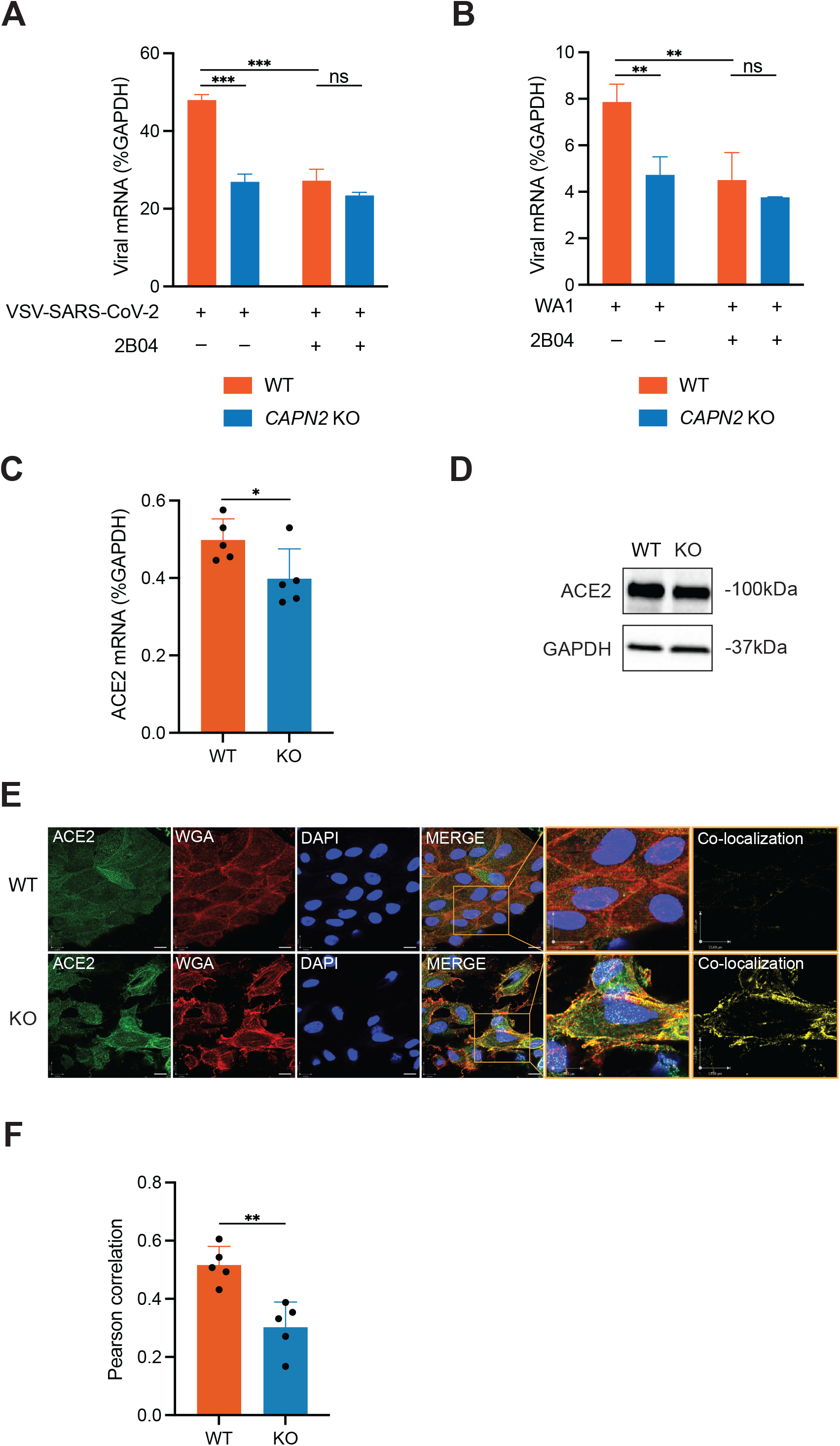
CAPN2 enhances ACE2 surface levels and spike-mediated virus attachment. (A) Cold binding assay with VSV-SARS-CoV-2. WT and *CAPN2* KO MA104 cells were infected by VSV-SARS-CoV-2 alone (MOI=20) or virus pre-incubated with a spike-targeted neutralizing antibody 2B04 (10 μg/mL) for 1 hour on ice followed by RNA extraction and RT-qPCR. Viral mRNA levels were measured and normalized to those of GAPDH. (B) Same as (A) except that SARS-CoV-2 strain WA1 was used instead. (C) ACE2 mRNA levels of WT and *CAPN2* KO cells. WT and *CAPN2* KO MA104 cells were harvested for RNA extraction followed by RT-qPCR analysis. The mRNA levels of ACE2 were measured and normalized to those of GAPDH. (D) Bulk ACE2 protein levels in WT and *CAPN2* KO cells. WT and *CAPN2* KO MA104 cells were harvested for western blot examining the levels of ACE2 and GAPDH. (E) Confocal analysis of surface levels of ACE2 in WT and *CAPN2* KO cells. WT and *CAPN2* KO MA104 cells were fixed and stained for surface glycoprotein (red, WGA), ACE2 (green), and nucleus (blue, DAPI). Scale bars: 13 μm. (F) Quantification of co-localization of ACE2 and WGA-stained cell membrane glycoprotein in WT and *CAPN2* KO MA104 cells.

To dissect the mechanism underlying reduced viral binding in the absence of CAPN2, we tested potential spike cleavage by CAPN2 given its role as a protease. We co-transfected HEK293 cells stably expressing human ACE2 with spike derived from WA1 strain, along with EGFP (control), CAPN2, transmembrane serine protease 2 (TMPRSS2), or furin, the major proteases known to cleave spike for efficient entry [16]. The cells lysates were harvested and the intensities of the full-length spike and its cleaved product S2 fragment were quantified (**Fig. S4A**). The spike cleavage efficiency was plotted as percentage of the cleavage product within overall spike protein levels. In this assay, overexpression of V5-tagged CAPN2 did not lead to significant spike cleavage more than EGFP, the negative control, when compared to other host proteases such as TMPRSS2 and furin (**Fig. S4B**).

Next, we examined the levels of SARS-CoV-2 cellular receptor ACE2 in WT and *CAPN2* KO cells. Interestingly, although the transcriptional level of ACE2 was slightly but statistically lower in *CAPN2* KO cells (**Fig. 5C**), this difference was not reflected on the bulk protein level (**Fig. 5D**). Of note, when we stained for subcellular localization of ACE2 in WT and KO cells, we observed much higher levels of surface ACE2 in WT cells co-localizing with wheat germ agglutinin (WGA) at the plasma membrane by confocal microscopy, in contrast to higher levels of intracellular ACE2 seen in the KO cells (**Fig. 5E**). Co-localization analysis of ACE2 and WGA, indicated by yellow signals in the inset images, showed that the surface ACE2 levels were significantly reduced in the *CAPN2* KO cells (**Fig. 5E and F**). Collectively, these data suggest that CAPN2 positively regulates the presence of ACE2 at the cell surface, thus enhancing spike-mediated SARS-CoV-2 binding and viral infectivity.

## DISCUSSION

SARS-CoV-2, like SARS-CoV, employs the spike protein that engages surface ACE2 to bind to host cells and is primed by TMPRSS2 and TMPRSS4 [17], as well as host cysteine proteases cathepsins B and L for entry into host cells [18]. Many proteases contribute to viral entry of SARS-CoV-2 and the development of immunopathology during COVID-19 diseases [17-21]. In this study, we uncovered the host protease CAPN2 as a novel host factor that aides the infection of SARS-CoV-2.

CAPN2 plays a major role in cancer-related cell proliferation [22, 23]. Although participation of calpains in virus infections has not been well understood, several published studies indicate pro-viral functions of the host gene CAPN2. CAPN2 expression was discovered to be an indicator of level of hepatic fibrosis during hepatitis B virus infection [24]. Additionally, CAPN2 enhances replication of echovirus 1 at a late stage step during the viral replication cycle [25] and CAPN2 promotes coxsackievirus entry into host cells [26]. In our study, we utilized the recombinant VSV-SARS-CoV-2 as a surrogate for SARS-CoV-2 and observed viral inhibition by calpain inhibitors through a series of experiments using chemical inhibitors, genetic knockouts, and classical virological approaches. Unlikethe literature describing the direct role of CAPN2 in viral entry and assembly, we show that CAPN2 also promotes SARS-CoV-2 infection by acting early to aid virus binding (**Fig. 5**). Further studies are needed to understand whether CAPN2 modulates ACE2 endosomal trafficking, recycling, or degradation.

Our current study has a number of limitations. The calpain inhibitors are not tested in primary human airway epithelial cells and their therapeutic utility is not yet explored to inhibit SARS-CoV-2 infection in relevant animal models. Another caveat is the frequency of mutations in the spike protein in SARS-CoV-2 strains. The relevance of CAPN2 to SARS-CoV-2 infection seems to be strain-specific (**Fig. 4C and D**), which we do not yet fully understand. Nonetheless, our findings highlight a novel function of CAPN2 in mediating SARS-CoV-2 entry and offer an alternative explanation to the protective efficacy of calpain inhibitors independent of blocking M^pro^ activities.

## MATERIALS AND METHODS

### Reagents, cells, and viruses

#### Reagents

MG132 (Selleckchem, S2619), gefitinib (Selleckchem, S1025), nigericin (InvivoGen, tlrl-nig/NIG-36-01), brefeldin A (Cell Signaling Technology, 9972S), FTY720 (Santa Cruz Biotechnology, sc-202161A), IBMX, concanamycin A (Enzo Life Sciences, ALX-380-034-C025), tetrandrine (Selleckchem, S2403), U18666A (Cayman Chemical, 10009085), ETP-46464 (Selleckchem, S8050), JIB-04 (Tocris, 4972), nitazoxanide (COVID Box, MMV688991), ketoconazole (COVID Box, MMV637533), AG-1478 (Selleckchem, S2728), caffeic acid (Selleckchem, S7414), thapsigargin (Cell Signaling Technology, 1278S), staurosporine (Cell Signaling Technology, 9953S), arbidol-HCl (Selleckchem, S2120). Calpain Inhibitor set includes ALLN, calpain inhibitor III, calpeptin, and E-64d used in the viral inhibition assays (208733-1SET, Sigma-Aldrich).

#### Cells

MA104 cells (ATCC, CRL-2378.1) were cultured in M199 medium (Thermo Scientific, 11150067) supplemented with 10% fetal bovine serum (FBS) and 1X Penicillin-Streptomycin-Glutamine (Thermo Scientific, 10378016). *CAPN2* KO MA104 cells were cultured in complete M199 medium with the addition of puromycin (10 μg/mL) for selection (single-guide RNA sequence: TGATCCGCATCCGAAATCCC). Vero E6 cells were cultured in DMEM (Thermo Scientific, 11965118) supplemented with 10% fetal bovine serum (FBS) and 1X Penicillin-Streptomycin-Glutamine.

#### Viruses

Recombinant VSV-eGFP [27] and VSV-eGFP-SARS-CoV-2 were previously described [8]. WT SARS-CoV-2 clone of the 2019n-CoV/USA_WA1/2020 (WA1/2020) strain and SARS-CoV-2 containing three point mutations in the spike gene E484K, N501Y, D614G were obtained from Pei-Yong Shi lab [28, 29], viruses stocks were propagated in stable clonal Vero-TMPRSS2 obtained from Sean Whelan lab.

#### Inhibitor Screen

Cells were seeded in 96 well plates. When they reached 80∼90% confluency, they were pretreated with indicated compounds at desired concentrations for 1 hour, followed by virus infection for 24 hours with the compound present. The cells were then washed and placed in clear PBS for Typhoon imager scanning. Co-encoded GFP serves as an indicator of infection level as the imager detects fluorescent signals. Darker color corresponds to more intense signals and therefore higher level of infection. The typhoon images were then processed using ImageJ for quantification of infection level.

#### Cell cytotoxicity assay

The cytotoxicity level of calpain inhibitors were determined using the Cell Counting Kit 8 (Abcam, ab228524). Cells in 96-well plates were treated with inhibitors of interest at concentrations within a range from 0.1 to 300 μM at 37°C for 25 hours. Fresh medium containing 10 μL of WST-8 substrate were added to each well to replace the inhibitor-containing medium. After 2 hours incubation at 37°C protected from the light, absorbance at 460nm was measured by BioTek ELx800 Microplate Reader and processed by Gen5 software.

#### Plaque Assay

MA104 cells were plated, grew to confluency in 6-well plates, and were infected with serial diluted viruses in serum-free M199 medium at 37°C for 1 hour. Afterwards, virus inoculum was replaced with warm agarose mixed with 2X M199 at 1:1 ratio. At 72 hours post infection, GFP signals in the plates were scanned by Amershad Typhoon 5 (GE) and plaque sizes were quantified by the ECHO microscope [27].

#### RNA extraction and quantitative PCR

RNA extraction were performed using QIAGEN RNeasy Mini kit (QIAGEN, 74104) per manufacturer’s instructions. For WT SARS-CoV-2 and triple variant E484K, N501Y, D614G infections, viral RNA was extracted using TRIzol (Invitrogen, 15596018) and chloroform following the product protocol. Viral mRNA levels (VSV N forward primer: 5’-GATAGTACCGGAGGATTGACGACTA-3’, VSV-N reverse primer: 5’-TCAAACCATCCGAGCCATTC-3’, SARS-CoV-2 N primer 1: 5’-ATGCTGCAATCGTGCTACAA-3’, primer 2: 5’-GACTGCCGCCTCTGCTC-3’, probe: 5’-/FAM/TCAAGGAACAACATTGCCAA/TAMRA/-3’) were examined by real-time RT-PCR using High Capacity cDNA Reverse Transcription kit (Applied Biosystems, 4368813) and AriaMX (Agilent) with 12.5 μl of either SYBR Green master mix (Applied Biosystems, 4367659) or Taqman master mix (Applied Biosystems, 4444557), reaching a total reaction volume of 25 μl. Expression of each gene was normalized to the expression of housekeeping gene GAPDH as previously described [30].

#### Western Blot

Cells were washed with PBS and lysed by RIPA buffer (Thermo Scientific, 89901) supplemented with 100X protease inhibitor cocktail and phosphatase inhibitor (Thermo Scientific, 78420), followed by a 10-minute incubation on ice. Cell lysates were then subjected to centrifugation at 13,500 RPM for 10 minutes at 4°C to remove cell debris and chromatids. The protein samples were then boiled in 2X Laemmli Sample Buffer (Bio-Rad, #1610737EDU) containing 5% β-mercaptoethanol at 95 °C for 5 minutes. Prepared samples were run in 4-12% gels and transferred onto nitrocellulose membranes. Membranes were blocked in 5% BSA in TBS + 0.1% Tween-20 (TBST) at room temperature before incubation at 4°C overnight with primary antibodies: SARS-CoV-2 spike RBD (Sino Biological, 40592-T62), GAPDH (BioLegend, 631402), calpain-2 (Cell Signaling Technology, 2539), ACE2 (R&D Systems, MAB933), S2 (Sino Biological, 40590-T62), and V5 (Cell Signaling Technology, 13202S). Membranes were then washed three times with TBST and incubated in secondary antibodies accordingly: anti-mouse HRP-linked IgG (Cell Signaling Technology, 7076S) or anti-rabbit HRP-linked IgG (Invitrogen, A27036) diluted in 5% BSA in TBST at room temperature for 1 hour. After the secondary antibody incubation, the membranes were washed three times with TBST and visualized by using Chemi-Doc imaging system (Bio-Rad).

#### Confocal microscopy

MA104 cells were seeded in eight-well chamber slides (catalog info here) and were fixed when reached 80% confluency in 4% paraformaldehyde for 10 min at room temperature. Cells were then washed with PBS once and stained with WGA (Thermo Scientific, W11262) for 10 minutes at room temperature. After another wash with PBS, cells were incubated with anti-ACE2 (Sino Biological, 10108-RP01-100) or isotype control (Cell Signaling Technology, 7074S) at room temperature for one hour. Stained cells were then washed with PBS once and then incubated with the secondary antibody (Invitrogen, A-11008) in dark for another hour. Postsecondary antibody incubation, the cells were washed and stained with DAPI (Invitrogen, P36962). The imaging was performed by a Zeiss LSM880 Confocal Microscope at the Molecular Microbiology imaging core facility at Washington University in St. Louis. Images were analyzed by Velocity v6.3 to generate co-localization and calculate the Pearson correlation coefficients.

#### Cold binding assay

Cells were seeded in 24-well plates and were ready for use when reached 60%∼80% confluency. Plates were pre-chilled on ice for 2∼4 hours prior to incubation with VSV-SARS-CoV-2 or SARS-CoV-2. An MOI of 20 was used to ensure maximum viral adsorption. Viruses, mixture of virus with 2B04, a neutralizing antibody against SARS-CoV-2 [15] were incubated at 37 °C for 1 hour. Pre-incubated virus, mixture of virus and antibody were chilled on ice for 30 minutes before added onto pre-chilled cells and incubated on ice. At 1 hour post incubation, the cells were washed with pre-chilled PBS three times and then lysed with RLT buffer for RNA harvesting.

#### Statistical analysis

Bar graphs are displayed as means ± SEM. Statistical tests were performed using GraphPad Prism 9.3.1. For **Figures 3B, 3C, 4A, 4C, 4D, 5C, and 5F**, statistical significance was calculated by Mann Whitney U test. For **Figure 5A and 5B**, statistical significance was calculated by two-way ANOVA Šidák’s multiple comparisons test. For **Supplemental Figure 4B**, statistical significance was calculated by one-way ANOVA Dunnett’s multiple comparisons test. For inhibition and cytotoxicity curves, EC_50_ and CC_50_ values in **Figures 1, 2 and Supplemental Figure 1** were calculated using nonlinear regression (curve fit). Asterisks indicate the following: *P≤0.05, **P≤0.01, and ***P≤0.001.

## Acknowledgement

We appreciate the helpful discussion of our studies with Dr. Haitao Guo (University of Pittsburgh), Dr. Jun Wang (Rutgers University), and Dr. Fumihiko Urano (Washington University in St. Louis).

## FIGURE LEGENDS

**Supplemental Figure 1. Nitazoxanide effectively inhibits SARS-CoV-2 infection**

(A) Inhibition and cytotoxicity of nitazoxanide (NTZ) against SARS-CoV-2-mNeonGreen infection. Vero E6 cells were treated with NTZ for 1 h prior to SARS-CoV-2-mNeonGreen infection at an MOI of 0.5. Infection level at 24 hpi was quantified based on immunofluorescence. For cytotoxicity measurement, cells were treated with NTZ at 0.1 μM to 1000 μM for 25 h before being subjected to WST-8 assay to test cell viability. Percentage cell cytotoxicity was plotted as a function of compound dosage. Both assays were repeated three times. EC_50_, CC_50_ and the SI (selectivity index) are as indicated.

(B) Same as (A) except VSV-SARS-CoV-2 and MA104 cells were used instead.

(C) Drug combination dose-response matrix and VSV-SARS-CoV-2 replication. MA104 cells were treated with NTZ along with JIB-04 at indicated concentrations for 1 h prior to infection by VSV-SARS-CoV-2 at an MOI of 3. GFP signals at 24 hpi were quantified to calculate the percentage of inhibition. Percent inhibition was plotted corresponding to color intensity.

**Supplemental Figure 2. VSV-SARS-CoV-2 infection is reduced in *CAPN2* KO cells**.

(A) GFP scans of WT and *CAPN2* KO MA104 cells infected by VSV-SARS-CoV-2 at indicated MOIs from 0.001 to 1, scanned at 6 to 48 hours post infection. The black dots represent GFP signals corresponding to infection levels.

(B) Viral mRNA levels in WT and *CAPN2* KO cells upon infection of VSV-SARS-CoV-2 at MOIs of 0.01 and 0.001, harvested at 24 and 48 hpi, respectively. The experiment was repeated twice.

**Supplemental Figure 3. A single clone of *CAPN2* KO MA104 cells was validated by Sanger sequencing**

sgRNA-targeted exon 7 of the *CAPN2* gene locus in WT (ref) and generated single clone CAPN2 KO (seq) by Sanger sequencing. Conserved and mutated regions were analyzed by CRISPR ID (http://crispid.gbiomed.kuleuven.be/).

**Supplemental Figure 4. WA1 spike is not cleaved significantly by CAPN2**

(A) WA1 spike cleavage by CAPN2, TMPRSS2, and furin shown by western blotting.

HEK293 cells were co-transfected with 0.5 μg of WA1 spike with 0.5 μg of EGFP, CAPN2, TMPRSS2, and furin, respectively. Protein samples were harvested at 24 hours post transfection. The result is representative of four repeats.

(B) WA1 spike cleavage quantification from 4 repeated experiments described above. Intensities of bands of full-length spike and cleaved product S2 were quantified using ImageJ.

