## Supplementary figures and images for "Calpain-2 mediates SARS-CoV-2 entry and represents a therapeutic target"

### Supplemental figures

#### Supplemental figure 1

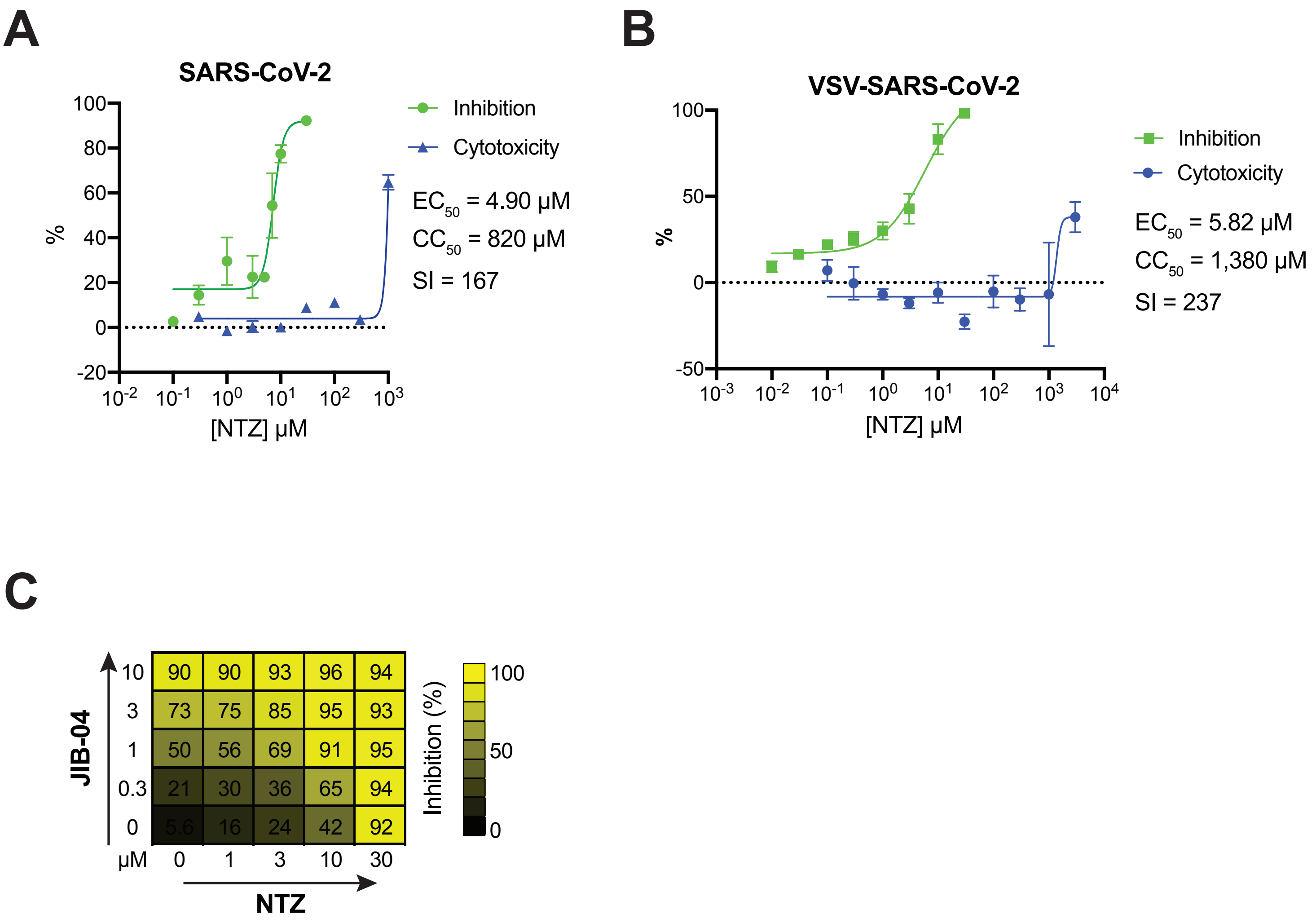

#### Supplemental figure 2

A

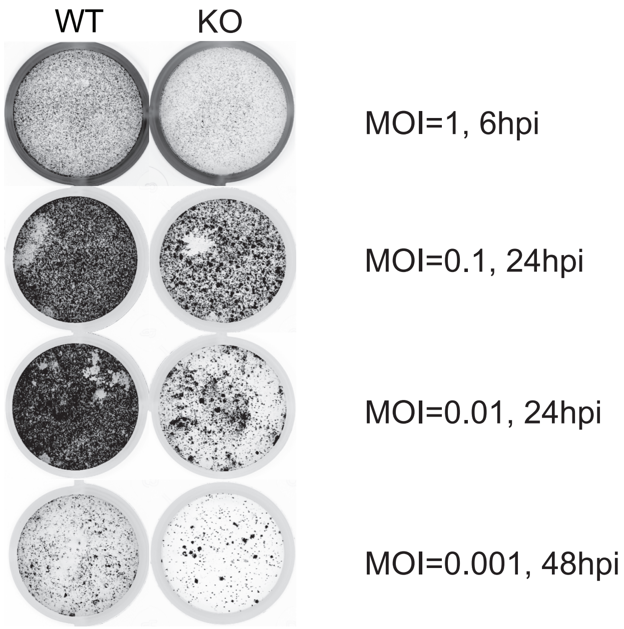


B

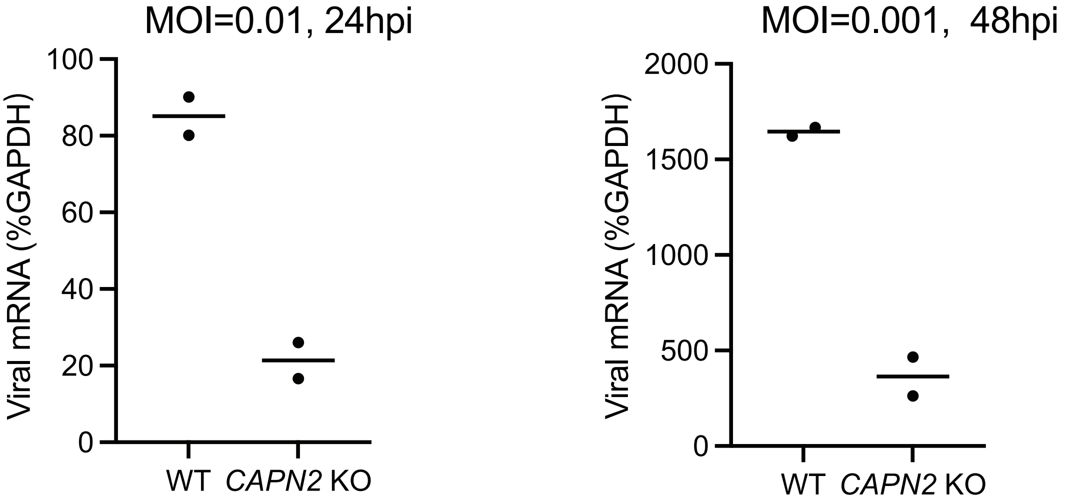

##### Supplemental figure 3

**A**

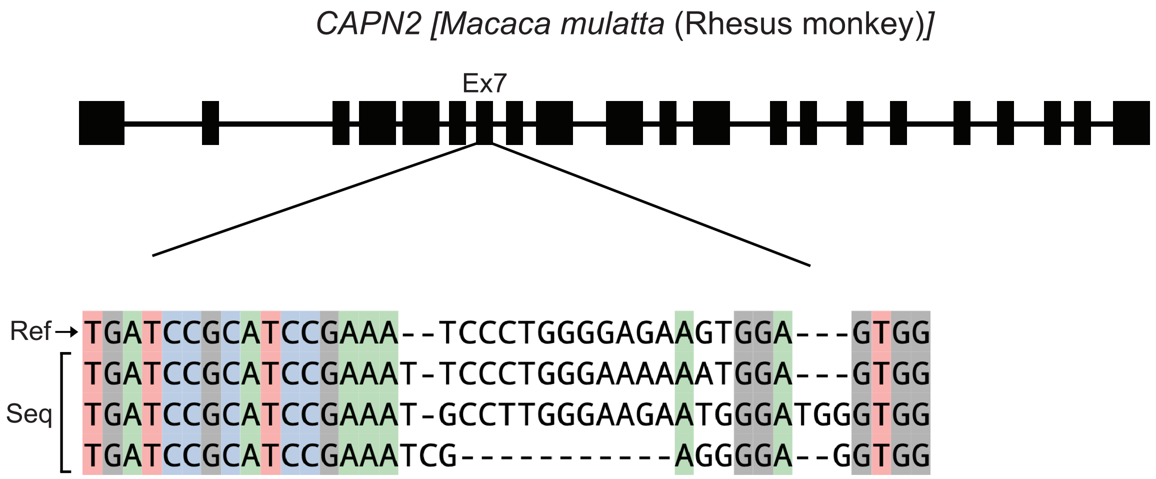

#### Supplemental figure 4

A

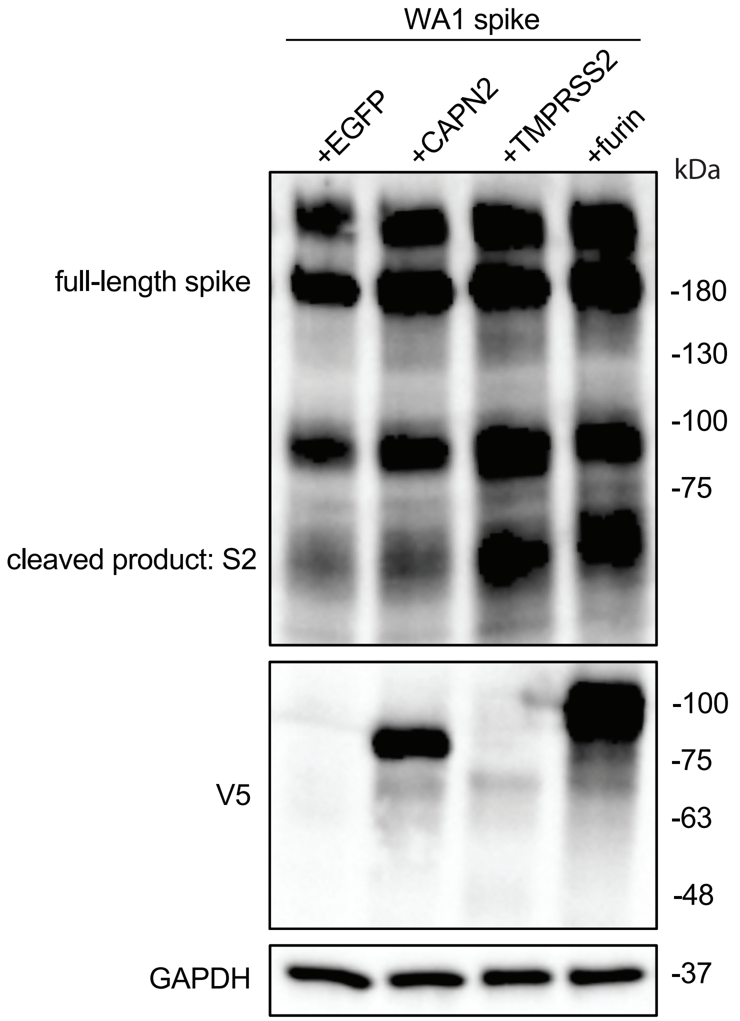


B

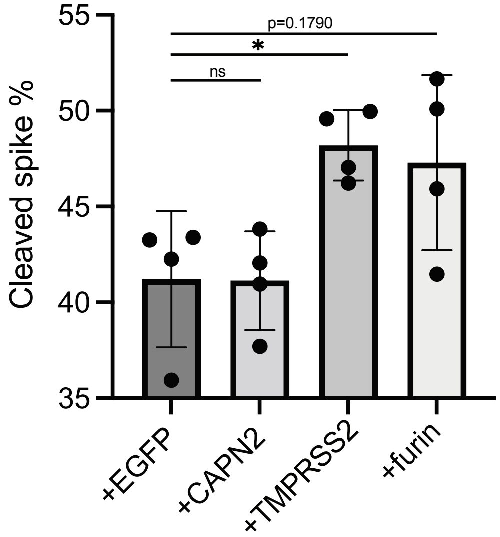
